# Deep learning for population genetic inference

**DOI:** 10.1101/028175

**Authors:** Sara Sheehan, Yun S. Song

## Abstract

Given genomic variation data from multiple individuals, computing the likelihood of complex population genetic models is often infeasible. To circumvent this problem, we introduce here a novel likelihood-free inference framework by applying deep learning, a powerful modern technique in machine learning. In contrast to Approximate Bayesian Computation, another likelihood-free approach widely used in population genetics and other fields, deep learning does not require a distance function on summary statistics or a rejection step, and it is robust to the addition of uninformative statistics. To demonstrate that deep learning can be effectively employed to estimate population genetic parameters and learn informative features of data, we focus on the challenging problem of jointly inferring natural selection and demography (in the form of a population size change history). Our method is able to separate the global nature of demography from the local nature of selection, without sequential steps for these two factors. Studying demography and selection jointly is motivated by *Drosophila,* where pervasive selection confounds demographic analysis. We apply our method to 197 African *Drosophila melanogaster* genomes from Zambia to infer both their overall demography, and regions of their genome under selection. We find many regions of the genome that have experienced hard sweeps, and fewer under selection on standing variation (soft sweep) or balancing selection. Interestingly, we find that soft sweeps and balancing selection occur more frequently closer to the centromere of each chromosome. In addition, our demographic inference suggests that previously estimated bottlenecks for African *Drosophila melanogaster* are too extreme, likely due in part to the unaccounted impact of selection.

## Introduction

With the advent of large-scale whole-genome variation data, population geneticists are currently interested in considering increasingly more complex models. However, statistical inference in this setting is a challenging task, as computing the likelihood of a complex population genetic model is a difficult problem both theoretically and computationally.

Approximate Bayesian Computation (ABC) [4,48] is a likelihood-free inference method based on simulating data and comparing their summary statistics. (A more detailed description of the framework is provided below.) This approach has been used to study various complex population genetic models (e.g., [5,27,47]) for which likelihood computation is prohibitive. Partly due to several influential theoretical works [3, 6,10,15, 29, 39, 44, 57], the popularity of ABC has grown rapidly over the past decade. ABC’s main advantages are that it is easy to use and is able to output a posterior distribution. There are a few challenges, however: 1) ABC uses a rejection algorithm, so the simulated data are not used optimally. 2) As such, training an ABC method requires a very large amount of simulated data. 3) The choice of a distance metric on summary statistics is an important consideration in designing an efficient ABC algorithm. 4) ABC suffers from the “curse of dimensionality,” with decreasing accuracy and stability as the number of summary statistics grows [2].

In this paper, we introduce an alternate likelihood-free inference framework for population genomics by applying *deep learning,* which is an active area of machine learning research. To our knowledge, deep learning has not been employed in population genomics before. A recent survey article [28] provides an accessible introduction to deep learning, and we provide a high-level description below. Our general goal in this paper is to demonstrate that statistical methods based on deep learning can allow more accurate inference of complex models than previously possible. As a concrete example, we consider models of non-equilibrium demography and natural selection, for which multi-locus full-likelihood computation is prohibitive.

To our knowledge, ABC has not been previously applied to the challenging problem of jointly inferring demography and selection. Several machine learning methods have been developed for selection, but they mostly focus on classifying the genome into neutral versus selected regions. Examples include methods based on support vector machines (SVMs) [45,46,51] or boosting [36,37,49]. Often these methods demonstrate robustness to different demographic scenarios, but do not explicitly infer demography.

The demographic models considered in this paper are restricted to a single population with time-varying effective population size, but the overall framework presented here can be applied to more general demographic models. Many methods have been developed to infer ancestral population size changes, including PSMC [34], diCal [56,58], and MSMC [53]. These methods assume that selection would not significantly bias the results. This is perhaps true for humans, but would not be true for *Drosophila,* for which selection seems rather ubiquitous throughout the genome.

A few previous works have addressed both population size changes and selection. Galtier *et al.* [16] developed a likelihood-based method for distinguishing between a bottleneck and selection. They applied their method to *Drosophila* data to conclude that a series of selective sweeps was more likely than a bottleneck, but did not explicitly infer parameters for both selection and demography. In their method, Galtier *et al.* assumed that demographic events affect the entire genome, whereas selection is a local phenomenon. In contrast, Gossmann *et al.* [20] estimated the effective population size locally along the genome, and reported that it is correlated with the density of selected sites. To make our results as easily interpretable as evolutionary events as possible, we estimate global effective population size changes.

To test our method, we simulate data under a variety of realistic demographies for *Drosophila melanogaster.* For each demography, we then simulate many regions with different selection parameters. We then apply our tailored deep learning method using a large number of potentially informative summary statistics. We demonstrate that parameters can be learned more accurately with deep learning than with ABC, in addition to being able to interpret which statistics are making the biggest contributions. After training our deep network, we also apply it to African *Drosophila, melanogaster* data to learn about its effective population size change history and selective landscape.

## ABC background

Rejection-based methods have been used since the late 1990’s (see [48,60]) to estimate population genetic parameters when the likelihood is difficult to compute. Early improvements to ABC quickly helped make it a popular method for a variety of scenarios (see [4] for a good introduction). ABC works by simulating many datasets under a prior for the parameters of interest. Then these datasets are reduced to a vector of summary statistics that are ideally informative for the parameters. The summary statistics that are closest to the summary statistics for the target dataset are retained, and the corresponding parameters used to estimate the desired posterior distributions. The definition of “close” is usually determined by the Euclidean distance on the summary statistic vectors, which can create biases if statistics are not properly normalized or have different variances.

Another problem with ABC is its inability to handle uninformative or weakly informative summary statistics. Intuitively, this is because these statistics add noise to the distance between two datasets; two datasets simulated under similar parameters may have some uninformative statistics that are far apart, or two datasets simulated under very different parameters may have some uninformative statistics that are close together. The distance can be further biased by differences in the magnitude of the summary statistics. Another major problem with ABC is the rejection step, which does not make optimal use of the datasets which are not retained. The more statistics and parameters used, the more datasets must be simulated and rejected to properly explore the space, making the interaction between these two issues even more problematic. One final issue with ABC is the black-box nature of the output. Given the distances between the simulated datasets and the target dataset, and the posterior, there is no clear way to tell which statistics were the most informative.

To tackle the problem of adding summary statistics, many methods for dimensionality reduction or selecting summary statistics wisely have been proposed (see [1,15,29,44], and [7] for a dimensionality reduction comparison). However, simple reductions cannot always learn subtle relationships between the data and the parameters. Expert pruning of statistics helps some methods, but given the lack of sufficient statistics, valuable information can be eliminated, especially when trying to infer many parameters. Blum and Francois [6] suggested performing a dimensionality reduction step on the summary statistics via a neural network similar to Fig. 1. This reduction is similar in spirit to the work presented here, although there are many algorithmic and application differences.

**Figure 1.**
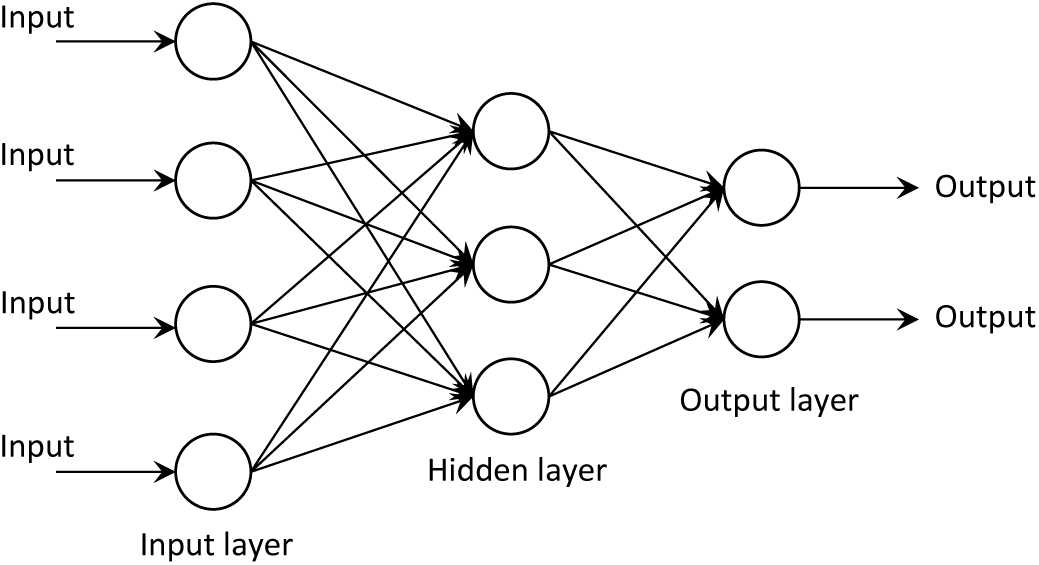
An example of a classical neural network. The single hidden layer serves to learn informative combinations of the inputs, remove correlations, and typically reduce the dimension of the data. After the optimal weight on each connecting arrow is learned through labeled training data, unlabeled data can be fed through the network to estimate the output parameters.

To address the problem of rejecting datasets, different weighting approaches have been proposed (see [6] for a good example of how the estimation error changes as fewer datasets are rejected). The idea is to keep more datasets, but then weight each retained dataset by the its distance to the target dataset. However, few approaches utilize all the datasets in this way, and the most popular implementation of ABC (*ABCtoolbox,* [61]) typically still rejects most of the simulated datasets by default.

### A brief introduction to deep learning

Deep learning has its beginnings in neural networks, which were originally inspired by the way neurons are connected in the brain [25]. Neural networks have been studied for over 60 years and a huge body of literature exists on the topic. Neural networks are typically used to learn complex functions between input data and output parameters in the absence of a model. While standard regression and classification methods involve fitting linear combinations of *fixed* basis functions, a neural network tries to *learn* basis functions (usually non-linear) appropriate for the data. A neural network architecture consists of multiple layers of *computational units* (nodes), with connections between the layers but not between nodes within a layer. Within a layer, each node computes a transformation (usually non-linear) of the outputs from the previous layer. Illustrated in Fig. 1 is a simple feed-forward neural network with a single hidden layer. The phrase “deep learning” refers to algorithms for learning *deep* neural network architectures with many hidden layers.

The *universal approximation theorem* [8,26] states that any continuous function on compact subsets of 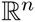 can be uniformly approximated by a feed-forward neural network with a single hidden layer, provided that the number of nodes in the hidden layer is sufficiently large and the transformation (called the activation function) associated with each node satisfies some mild conditions. However, it can be challenging to learn the weights of such a network and to interpret the hidden layer. So as learning problems became more complex, it was desirable to train networks with more hidden layers. Since their introduction over 30 years ago, deep architectures have proved adept at modeling multiple levels of abstraction. However, they were notoriously difficult to train since their objective functions are non-convex and highly non-linear, and the level of non-linearity increases with the number of layers in the network. A major breakthrough was made in 2006 when Hinton and Salakhutdinov [23] showed that a deep feed-forward neural network can be trained effectively by first performing unsupervised “pretraining” one layer at a time, followed by supervised fine-tuning using a gradient-descent algorithm called *back-propagation* [52]. (Simply put, pretraining provides a good initialization point for non-convex optimization.) They applied their learning algorithm to dimensionality reduction of images and achieved substantially better results than PCA-based methods.

Following the work of Hinton and Salakhutdinov, deep learning has been applied to various challenging problems in computer science over the last 5 years, making groundbreaking progress. Deep learning broke long-standing records for accuracy that had been set by approaches based on hand-coded rules. Well-known examples include automatic speech recognition (transforming spoken words into typed text) [21,40] and computer vision (automatically classifying images into different categories and tagging objects/individuals in photos) [31].

Many variations have been developed, including *dropout*, which attempts to learn better and more robust features of the data (see [9,24]). Deep learning has also been applied to problems in neuroscience [30] and computational biology [33,50,62], but has not been used for population genetics before. We demonstrate here that deep learning provides solutions to some of the problems with ABC, and can provide an alternative to traditional likelihood-free inference in population genetics. It could also be used to select optimal statistics for ABC or another method.

## Results

In this paper, we describe how we apply deep learning to the challenging problem of jointly estimating demography and selection (see [35] for a recent review of this topic). One reason why this problem is difficult is that demography (for example, a bottleneck in population size) and selection can leave similar signals in the genome. Untangling the two factors directly has rarely been attempted; most methods that estimate selection try to demonstrate robustness to demographic scenarios, rather than estimating demographic parameters jointly with selection. Our analysis is motivated by *Drosophila*, where previous demographic estimates may have been confounded by pervasive selection, such as the demographic history inferred by Duchen *et al.* [12]. The reverse has occurred as well, with selection estimates being confounded by demography [19]. See [55] for a more thorough discussion of the role selection plays in the *Drosophila* genome. In what follows, we first test the performance of our method on simulated data and then apply it to analyze 197 *Drosophila melanogaster* genomes from Zambia, Africa [32].

### Simulating data

To create a simulated dataset that is appropriate for our scenario of interest, we first define the parameters we would like to estimate. For simplicity, we consider piecewise-constant population size histories with three epochs: a recent population size *N*_1_, a bottleneck size *N_2_,* and an ancient size *N_3_.* Further, we define a genomic region as belonging to 4 different selection classes: no selection (neutral), positive directional selection (hard sweep), selection on standing variation (soft sweep), and balancing selection. See [13] for a more complete analysis of the different types of selection in *Drosophila.* To accurately reflect that demography affects the entire genome while selection affects a particular region, we simulate many genomic regions under the same demographic history, but the selection class for each region is chosen independently. See Fig. 2 for a simplified illustration of the data.

**Figure 2.**
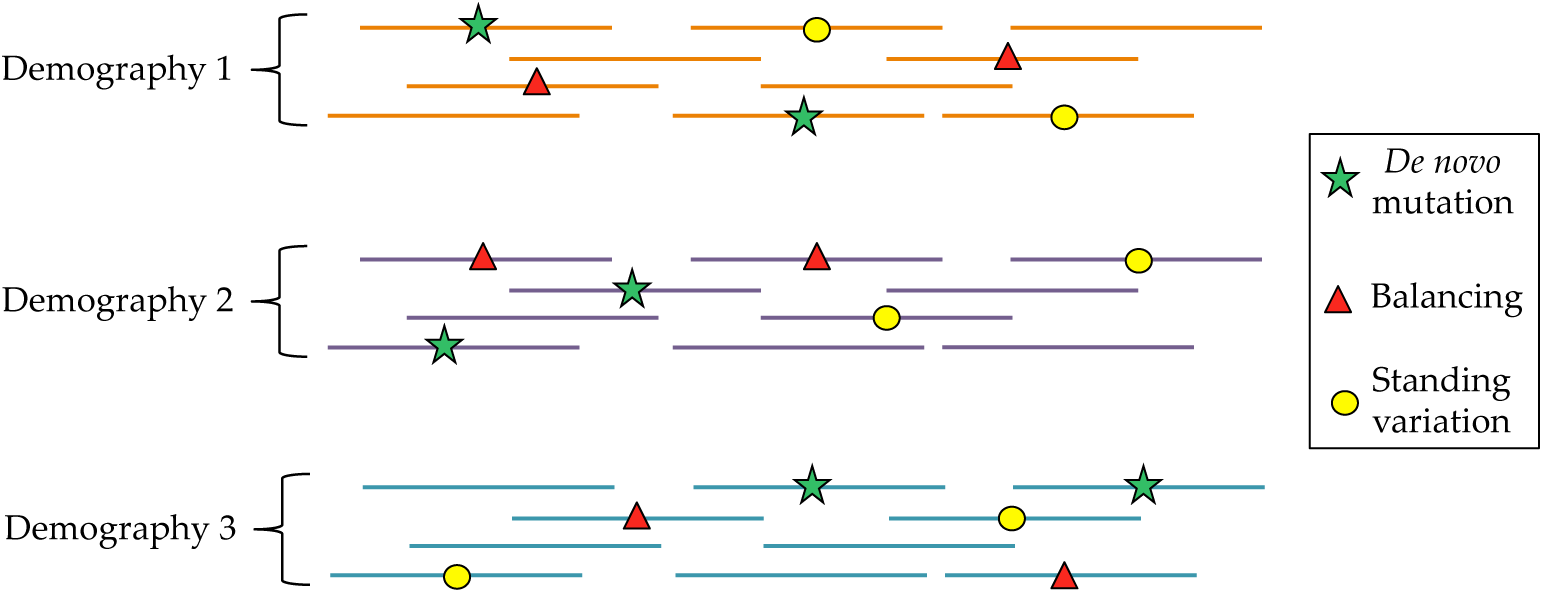
Input data for the demography and selection scenario. For each demographic history (bottleneck), we simulated many different genomic regions. Each region can either have no selection (neutral), one site with an *de novo* mutation under positive selection (hard sweep), one site under balancing selection, or one standing variant under positive selection (soft sweep).

To simulate data, we used the program `msms` [14]. To make our simulated data as close to the real data as possible, we simulated *n* = 100 haplotypes, and correspondingly downsampled the Zambia *Drosophila melanogaster* dataset to match. We repeated the following procedure 2500 times. First we selected three population sizes for the demographic model, then simulated 160 regions with these sizes, 40 for each selection scenario. Each region was 100 kb, with the selected site (if present) occurring randomly in the middle 20 kb of the region. We used a baseline effective population size *N*_ref_ = 100,000, a per-base, per-generation mutation rate *μ* = 8.4 × 10^−9^ [22], and a per-base, per-generation recombination rate *r* equal to *μ*, as inferred by PSMC [34]. We used a generation time of 10 generations per year. Based on the population size change results of PSMC, we chose the time change-points for demographic history to be *t*_1_ = 0.5 and *t*_2_ = 5 in coalescent units. Scaled effective population size parameters λ*_i_*:= *N_i_/N*_ref_ and their prior distributions are below:

1. Recent effective population size scaling factor: λ_1_ ~ Unif(3,14).
2. Bottleneck effective population size scaling factor: λ_2_ ~ Unif(0.5,6).
3. Ancient effective population size scaling factor: λ_3_ ~ Unif(2,10).

For the selection classes, the different types are shown below:

- Class **0, neutral**: no selection, neutral region.
- Class **1, hard sweep:** positive selection on a *de novo* mutation (i.e., hard sweep). For the selection coefficient, we used *s* ∈ {0.01,0.02,0.05,0.1}, with 10 regions for each value of s. The onset of selection was chosen to be 0.005 in coalescent units, which provided sweeps at a variety of different stages of completion at the present time. We discarded datasets where the frequency of the selected allele was 0 (early loss of the beneficial mutation due to drift), but for a large fraction of datasets, the selected allele had not yet fixed.
- **Class 2, soft sweep**: positive selection on standing variation (i.e., soft sweep). The selection coefficients and selection start time were chosen as in the hard sweep scenario, but now the initial frequency of the selected allele was chosen to be 0.001.
- **Class 3, balancing**: heterozygote advantage, balancing selection. The selection start time and selection coefficients were chosen in the same fashion as Class 1.

Using this strategy, we simulated 2500 different demographic histories, with 160 regions for each one, for a total of 400,000 datasets. To build 160 regions for each demography, we simulated 40 datasets for each of the classes above.

### Transforming input data into summary statistics

For many deep learning applications, the raw data can be used directly (the pixels of an image, for example). Unfortunately, we currently cannot input raw genomic data into a deep learning method. Similarly to ABC, we need to transform the data into summary statistics that are potentially informative about the parameters of interest. Unlike ABC, however, deep learning should not be negatively affected by correlated or uninformative summary statistics. Thus we sought to include a large number of potentially informative summary statistics of the data. To account for the impact of selection, we divided each 100 kb region into three smaller regions: 1) close to the selected site (40-60 kb), 2) mid-range from the selected site (20-40 kb and 60-80 kb), and 3) far from the selected site (0-20 kb and 80-100 kb). These regions are based off of the simulation scenario in Peter *et al.* [47], and shown more explicitly in Fig. 3. Within each of these three regions, we calculated the following statistics, except where noted.

**Figure 3.**
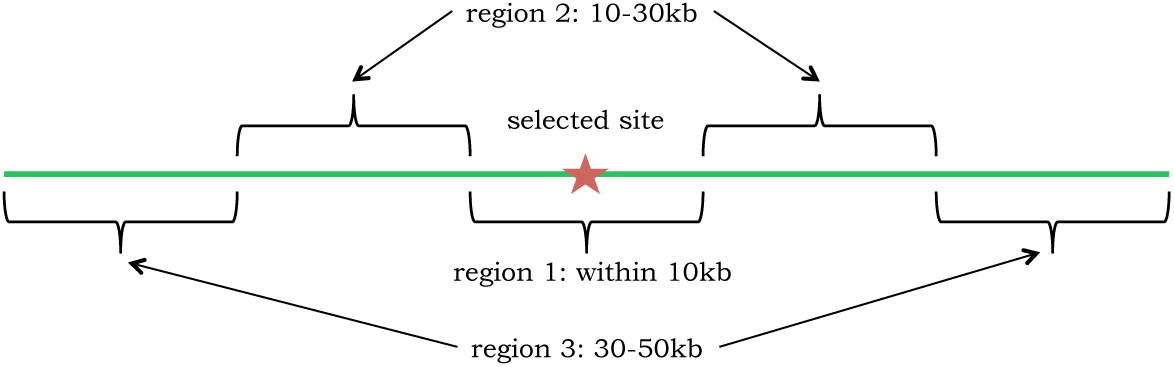
Regions used for computing the statistics, which are based off of Peter *et al.* [47]. Note that the selected site was chosen randomly within region 1.

For all the statistics described below, *n* is the haploid sample size. In the case of simulated data, *n* = 100. For the real data, we had 197 samples; within each 100 kb region we sorted the samples by missing data, then retained the 100 most complete samples, except where noted.

1. Number of segregating sites within each smaller region, *S*. Since all of the statistics must be in [0,1], we normalized by *S*_max_ = 5000 (any *S > S*_max_ was truncated): 3 statistics.
2. Tajima’s *D* statistic [59], computed as follows

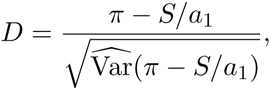

where π is the average number of pairwise differences between two samples, and 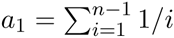. We normalized by *D*_min_ = −3.0 and *D*_max_ = 3.0, again truncating the rare statistic outside this range: 3 statistics.
3. Folded site frequency spectrum (SFS): *η_i_* is the number of segregating sites where the minor allele occurs *i* times out of *n* samples, for *i* = 1,2, …, ⌊n/2⌋. For the real data, for each segregating site, enough samples were included to obtain 100 with non-missing data. If that was not possible for a given site, the site was not included. To normalize the SFS, we divided each *η_i_* by the sum of the entries, which gives us the probability of observing *i* minor alleles, given the site was segregating: 50 · 3 = 150 statistics.
4. Length distribution between segregating sites: let *B_k_* be the number of bases between the *k* and *k* + 1 segregating sites. To compute the distribution of these lengths, we define *J* bins and count the number of lengths that fall into each bin:

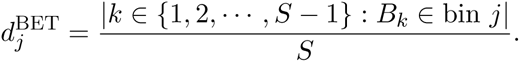 We choose *J* = 16 equally spaced bins, the first starting at 0 and the last starting at 300: 16 · 3 = 48 statistics.
5. Identity-by-state (IBS) tract length distribution: for each pair of samples, and IBS tract is a contiguous genomic region where the samples are identical at every base (delimited by bases where they differ). For all pairs, let *L* be the set of IBS tract lengths. In a similar fashion to the length distribution statistics, we define *M* bins, and count the number of IBS tract lengths that fall into each bin:

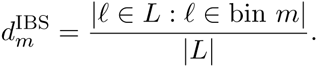 We choose *M* = 30 equally spaced bins, the first starting at 0 and the last starting at 5000: 30 · 3 = 90 statistics.
6. Linkage disequilibrium (LD) distribution: LD is a measure of the correlation between two segregating sites. For example, if there was no recombination between two sites, their alleles would be highly correlated and LD would be high in magnitude. If there was infinite recombination between two sites, they would be independent and have LD close to 0. For two loci, let *A, a* be the alleles for the first site, and *B,b* be the alleles for the second site. Let pa be the frequency of allele *A, p_B_* be the frequency of allele *B*, and *pab* be the frequency of haplotype *AB*. Then the linkage disequilibrium is computed by

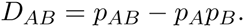 Here we compute the LD between pairs of sites, where one site is in the “selected” region, and the other is in each of the three regions (including the selected region). Then we create an LD distribution similar to the IBS distribution above, using 16 bins, with the first bin ending at −0.05, and the last bin starting at 0.2: 16 · 3 = 48 statistics.
7. H1, H12, and H2 statistics, as described in Garud *et al.* [17]. These statistics help to distinguish between hard and soft sweeps, and are calculated in the selected (middle) region only: 3 statistics.

This gives us a total of 345 statistics.

### Comparison with ABCtoolbox

First we wanted to compare the performance of deep learning with that of ABC. We used the popular ABCtoolbox [61], using the same training and testing datasets as for deep learning. However, since ABC is not well suited to classification, we restricted this analysis to estimating (continuous) demographic parameters only. For ABC, the training data represents the data simulated under the prior distributions (uniform in our case), and each test dataset was compared with the training data separately. We retained 5% of the training datasets, and used half of these retained datasets for posterior density estimation. Overall, we used 75% of the datasets for training and 25% for testing.

We tested two scenarios, one with the full set of summary statistics (345 total), and the other with a reduced set of summary statistics (100 total). For the reduced set of summary statistics, we chose statistics which seemed to be informative: the number of segregating sites, Tajima’s *D*, the first 15 entries of site frequency spectrum, H1, and the distribution of distances between segregating sites. The results are shown in Table 1, which suggest that deep learning produces more accurate estimates of the recent population size (*N*_1_) than does ABCtoolbox, whereas they have comparable accuracies for more distant past sizes.

**Table 1.** A comparison between ABCtoolbox and deep learning for demography only. Out of 1000 demographies (160,000 datasets total), 75% were used for training and 25% for testing. In this scenario, deep learning generally outperforms ABCtoolbox, as measured by the relative error: |N_est_ – *N*_true_ | /*N*_true_. There is generally more improvement using deep learning when the number of statistics is larger.

| Dataset | Method | $N_1$ error | $N_2$ error | $N_3$ error |
| --- | --- | --- | --- | --- |
| Full summary statistics | ABCtoolbox | 0.062 | 0.043 | 0.218 |
|  | Deep learning | 0.044 | 0.028 | 0.221 |
| Filtered summary statistics | ABCtoolbox | 0.161 | 0.035 | 0.311 |
|  | Deep learning | 0.065 | 0.055 | 0.319 |

### Results for demography and selection on simulated data

Moving to the demography and selection scenario, we used the dataset and summary statistics described above, which contains 400,000 datasets total. We used 75% for training and 25% for testing. In Table 2, we show the population size results for a network with 3 hidden layers of sizes 25, 25, and 10. The best results were found when we averaged the statistics for all of the datasets with the same demography (160 datasets, or “regions” each).

**Table 2.**
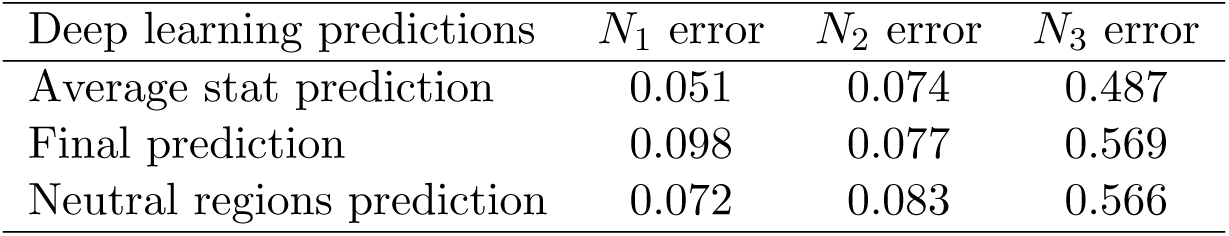
Deep learning results for population sizes under the scenario with demography and selection, using a network with 3 hidden layers of sizes 25, 25, and 10. We evaluate the results in three ways, using the relative error of the estimates: |*N*_est_ – *N*_true_|/*N*_true_. First, for each test demography, we average the statistics for each dataset, and then run these values through the training network. Second, for each test demography, we run the datasets through the network one by one, then average the predictions. Finally, we perform the second procedure, but with only the regions we classified as neutral. We note that the most ancient size (*N*_3_) is always the least accurately estimated.

| Deep learning predictions | $N_1$ error | $N_2$ error | $N_3$ error |
| --- | --- | --- | --- |
| Average stat prediction | 0.051 | 0.074 | 0.487 |
| Final prediction | 0.098 | 0.077 | 0.569 |
| Neutral regions prediction | 0.072 | 0.083 | 0.566 |

Table 3 shows the results for an example demography, with 95% confidence intervals (CI) calculated from the estimates for each (predicted) neutral region within the genome. In this example (and more generally), the most ancient size *N*_3_ is the least accurately estimated. When considering a single dataset, there is not always a clear winner among our three prediction methods. Overall, the average statistic method is usually the most accurate, followed by the neutral regions method.

**Table 3.**
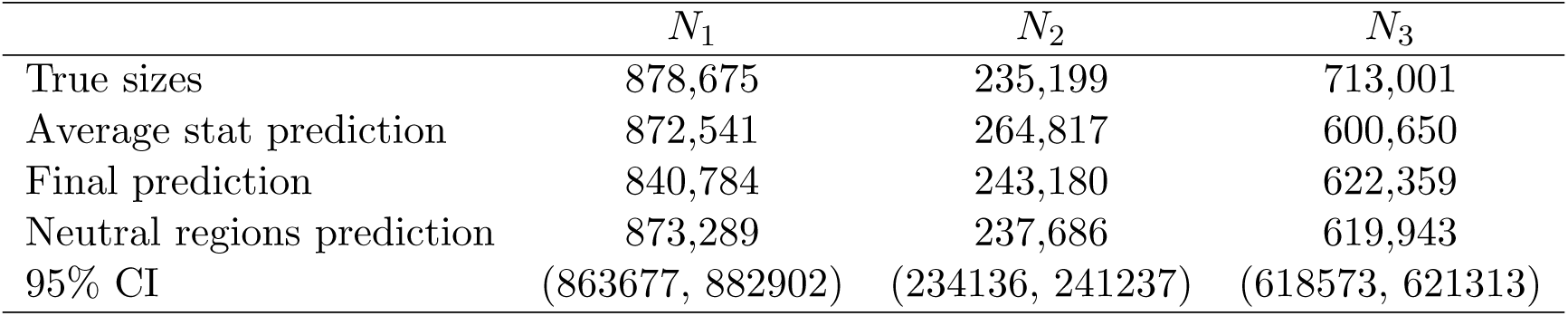
Results for an example demography, with the true population sizes shown in the first row. The next three rows represent predictions of the sizes, with the prediction based on neutral genomic regions being generally the most accurate. For this final prediction, 95% confidence intervals are shown in the last row.

To analyze the selection results, we calculated a confusion matrix in Table 4 to show which datasets of each class were classified correctly, or misclassified as belonging to a different class. Our most frequent error is classifying hard sweep datasets as neutral (row 2, column 1 of the confusion matrix). We hypothesized that this was either because selection occurred anciently and quickly, or because selection occurred recently and the sweep was not yet complete. To test this, we examined the results conditional on the frequency of the selected allele at present. The results shown in Table 5 suggest that in fact many of the sweeps were not complete, which is why the regions sometimes appeared neutral. Regardless, this type of false negative error is arguably preferable to falsely calling many regions to be under selection when they are in fact neutral.

**Table 4.** Confusion matrix for the selection predictions, in the demography and selection scenario. Each row represents the datasets that truly belong to each selection class. Each column represents the datasets that were actually classified as each selection class. Ideally we would like all 1’s down the diagonal, and 0’s in the off-diagonal entries. The largest number in each row is shown in boldface. We can see that neutral datasets are the easiest to classify, and sometimes regions under selection (hard sweeps in particular) look neutral as well (first column). The overall percentage of misclassified datasets was 6.2%.

| True Class | Called Class |  |  |  |
| --- | --- | --- | --- | --- |
|  | Neutral | Hard Sweep | Soft Sweep | Balancing |
| Neutral | **0.9995** | 0.0002 | 0.0003 | 0.0000 |
| Hard Sweep | 0.1434 | **0.8333** | 0.0032 | 0.0201 |
| Soft Sweep | 0.0096 | 0.0010 | **0.9891** | 0.0003 |
| Balancing | 0.0301 | 0.0356 | 0.0056 | **0.9287** |

**Table 5.** Hard sweep results, broken down by the present-day frequency (*f*) of the selected allele (which we would not know for a real dataset). We defined “low” frequency as *f* ∈ [0, 0.3), “moderate” frequency as *f* ∈ [0.3,0.7), and “high” frequency as *f* ∈ [0.7,1]. We can see that if the frequency of the selected allele is low, the region is often misclassified as neutral, since the selective sweep is not yet complete. However, if the frequency is moderate or high, the dataset is usually classified correctly as a hard sweep.

| $f$ | | Called Class | | | |
| --- | --- | --- | --- | --- | --- |
|  |  | Neutral | Hard Sweep | Soft Sweep | Balancing |
| Low | (18.4% of datasets) | **0.6820** | 0.3137 | 0.0009 | 0.0035 |
| Moderate | (12.5% of datasets) | 0.0988 | **0.8220** | 0.0000 | 0.0793 |
| High | (69.1% of datasets) | 0.0084 | **0.9734** | 0.0044 | 0.0138 |

We also wanted to test the impact of unsupervised pretraining using autoencoders (see `Methods`). Tables 6 and 7 compare the results for a randomly initialized network to a network initialized using autoencoders. These results demonstrate that pretraining is very effective. Due to the non-convex nature of the optimization problem, random initialization is most likely finding a poor local minima for the cost function.

**Table 6.** Relative error on the test dataset, for a deep network with 6 hidden layers. For the results in the first row, the weights of the entire network were initialized randomly, then optimized. In the second row, the weights were initialized using autoencoders for each layer. The positive impact of unsupervised pretraining is clear; random initialization causes the optimization to get stuck in a local minima.

| Initialization Type | $N_1$ error | $N_2$ error | $N_3$ error |
| --- | --- | --- | --- |
| Random | 0.429 | 0.421 | 0.710 |
| Autoencoder | 0.061 | 0.166 | 0.577 |

**Table 7.** Confusion matrix for the selection predictions, compared between random initialization (top) and autoencoder initialization (bottom), for a deep network with 6 hidden layers. Again, ideally we would like all 1’s down the diagonal, and 0’s in the off-diagonal entries. The largest number in each row is shown in boldface. When the network is initialized randomly, almost every dataset is classified as neutral; the network has not really learned anything meaningful from the input data. The overall percentage of misclassification is 74.8% for random initialization, while it is only 6.1% for autoencoder initialization.

| True Class | Called Class |  |  |  |
| --- | --- | --- | --- | --- |
|  | Neutral | Hard Sweep | Soft Sweep | Balancing |
| Random Initialization |  |  |  |  |
| Neutral | **1.000** | 0.000 | 0.000 | 0.000 |
| Hard Sweep | **0.978** | 0.007 | 0.000 | 0.015 |
| Soft Sweep | **1.000** | 0.000 | 0.000 | 0.000 |
| Balancing | **1.000** | 0.000 | 0.000 | 0.000 |
| Autoencoder Initialization |  |  |  |  |
| Neutral | **1.000** | 0.000 | 0.000 | 0.000 |
| Hard Sweep | 0.145 | **0.831** | 0.004 | 0.021 |
| Soft Sweep | 0.011 | 0.001 | **0.987** | 0.000 |
| Balancing | 0.030 | 0.028 | 0.001 | **0.941** |

### Results on real *Drosophila melanogaster* data

We then ran the real *Drosophila melanogaster* data through this trained network just like any other test dataset. We considered only the chromosome arms 2L, 2R, 3L, and 3R in this study. We partitioned each chromosome arm into 20 kb windows and ran our method on five consecutive windows at a time, sliding by 20 kb after each run. Classification was performed on the middle 20 kb window.

For the demography, the population size results are shown in Table 8. The first row (average statistic method) is our best estimate, which we also plot and compare with other histories in Fig. 4. Our history is close to the PSMC result, although more resolution would be needed for a proper comparison. The expansion after the bottleneck is roughly consistent with previous results citing the range expansion of *Drosophila melanogaster* (out of sub-Saharan Africa) as beginning around 15,000 years ago [18]. This range expansion likely led to an effective population size increase like the one we infer.

**Table 8.** Population size results for African *Drosophila* from Zambia, rounded to the nearest hundred. The first row (predictions based on averaging the statistics for each region in the *Drosophila* genome) represents our best estimate of the population sizes. The final row is a 95% confidence interval.

| Prediction | $N_1$ (recent) | $N_2$ (bottleneck) | $N_3$ (ancestral) |
| --- | --- | --- | --- |
| Average statistics | 544,200 | 145,300 | 652,700 |
| Average predictions | 622,100 | 158,200 | 650,400 |
| Neutral regions | 607,300 | 161,400 | 651,200 |
| 95% CI | (596,700 – 617,900) | (157,800 – 165,000) | (649,600 – 652,700) |

**Figure 4.**
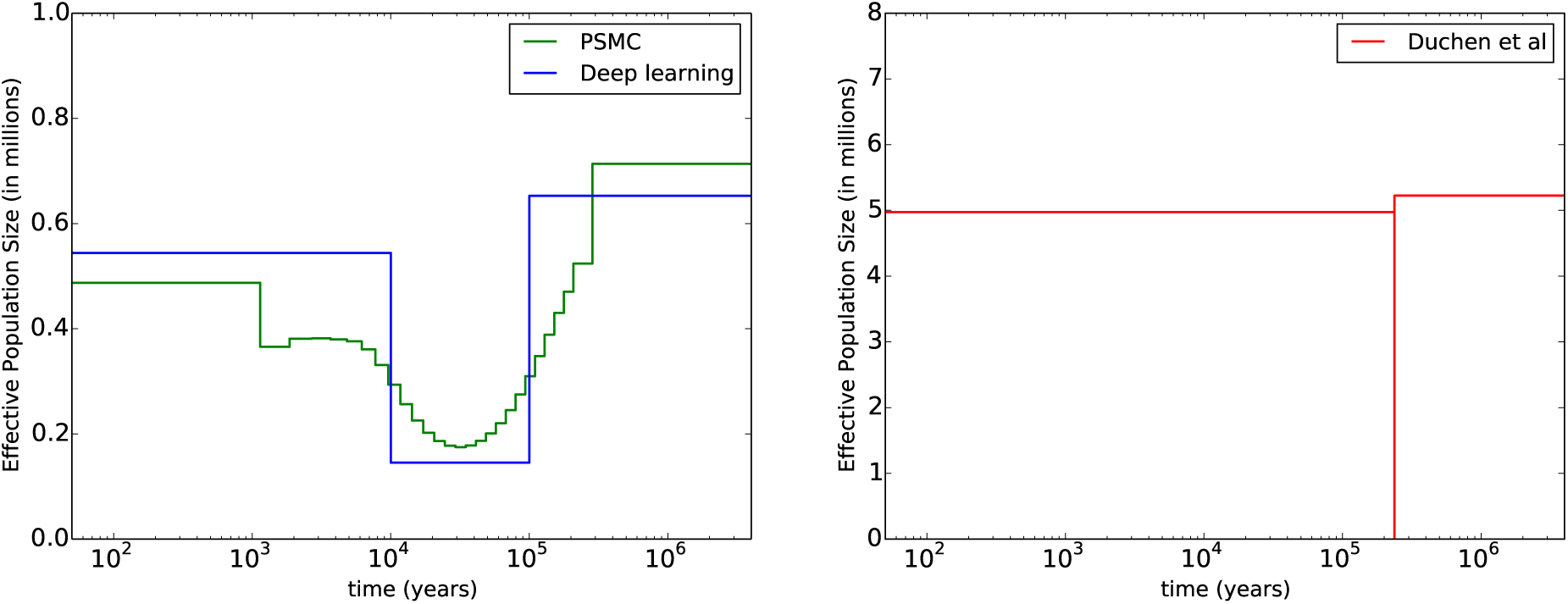
Comparison between demographic histories. On the left in green is PSMC [34], run on the entire genome for a subset of the data (*n* = 20). In blue is our history from the first line of Table 8. On the right in red is the history from Duchen *et al.* [12], which assumes a very short, severe bottleneck (note the change in the *y*-axis scale). A more gradual bottleneck seems more realistic, although we do not have a simple explanation for why there was a bottleneck at all.

The number of windows classified as Neutral, Hard Sweep, Soft Sweep, and Balancing Selection are 1191, 2572, 429, and 637, respectively. See Table S1 for further details. If we restrict our analysis to regions classified with probability greater than 0.9999, then we find 47 hard sweeps, 69 soft sweeps, and 18 regions under balancing selection. In Table S2, we include a table of these high-confidence windows, along with which genes are found in each window. We also include a plot (Figure S1) of where the selected regions fall throughout the genome (restricted to regions with a probability at least 0.95). Interestingly, soft sweeps and balancing selection seem to occur more frequently closer to the centromere of each chromosome. There is also an excess of hard sweeps on chromosome arm 2L.

Upon examining the genes in the top regions classified to be under selection, we find several notable results. In the hard sweep category, a few of the top regions harbor many genes involved in chromatin assembly or disassembly. Further, there are two regions each containing exactly one gene, where the gene has a known function. These are *Fic domain-containing protein* (detection of light and visual behavior), and *charybde* (negative regulation of growth). In the soft sweep category, we find many genes related to transcription, and several related to pheromone detection (chemosensory proteins). We also find four regions each containing exactly one gene, with that gene having a known function. These are *gooseberry* (segment polarity determination), *steppke* (positive regulation of growth), *Krüppel* (proper body formation), and *Accessory gland protein 36DE* (sperm storage, [41]). Fly embryos with a mutant *Krüppel* (German for “cripple”) gene have a “gap” phenotype with the middle body segments missing [38]. Finally in the balancing selection category, one interesting gene we find is *nervana 3,* which is involved in the sensory perception of sound. We also find balancing selection regions containing the genes *cycle* (circadian rhythms) and *Dihydropterin deaminase* (eye pigment), although there are other genes within these putatively selected regions.

We also wanted to investigate which statistics were the most informative for our parameters of interest, using the procedure described in `Methods` (c.f., Algorithm 1). To that affect, we kept the 25 most informative statistics for each population size and selection, highlighting common statistics in the 4-way Venn diagram in Fig. 5. For each statistic name in the diagram, *close, mid,* and *far* represent the genomic region where the statistic was calculated. The numbers after each colon refer to the position of the statistic within its distribution or order. For the SFS statistics, it is number of minor alleles. An interesting set of four statistics is informative for all parameters, including very long IBS tracts close to the selected site (“IBS, close: 30”) and the H12 statistic. Additionally, “LD, mid: 4” represents low LD between sites close to the selected site and sites mid-range from the selected site (likewise for LD “LD, far: 4”). Many low LD pairs could signal a lack of selection, and vice versa. IBS statistics are generally not as helpful for selection, but extremely informative for the population sizes, especially *N*_2_ and *N*_3_. Bottlenecks can have a significant impact on IBS tract distributions, so it makes sense that *N*_2_ (the bottleneck size) is the most reliant on IBS statistics. The number of segregating sites *S* is generally quite informative, especially for selection. Interesting, the folded site frequency spectrum (SFS) is not as informative as one might have anticipated. However, the number of singletons is very informative, especially for selection and *N*_1_, which could have been foreseen given that recent events shape the terminal branches of a genealogy, and thus the singletons. Tajima’s *D* is helpful for selection only, which it was designed to detect. The distances between segregating sites (“BET” statistics) do not generally seem very helpful. Neither does H2, the frequency of the second most common haplotype. It is comparing H1 to H2 (via the H12 statistic) that is helpful.

**Figure 5.**
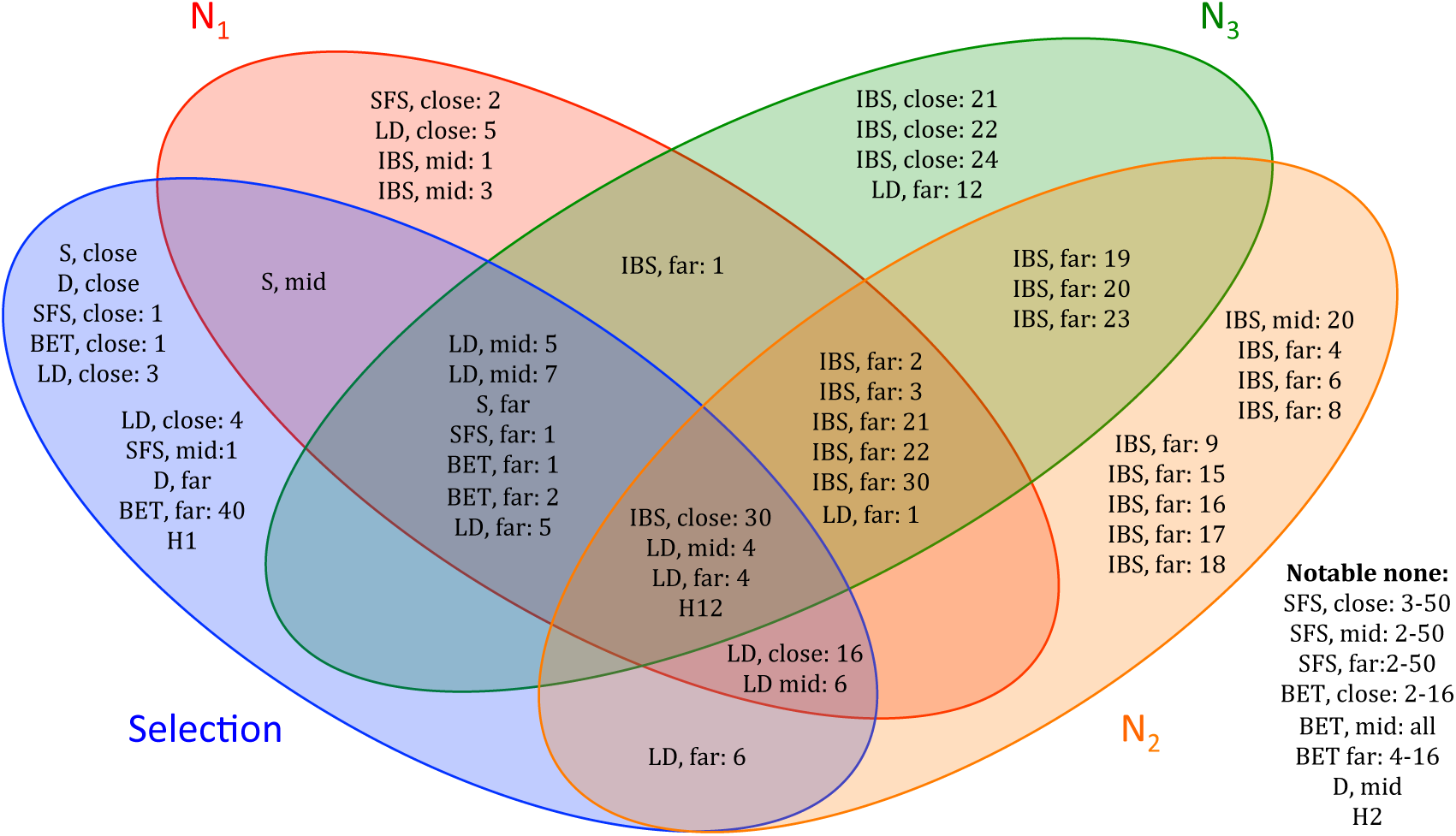
A Venn diagram of most informative statistics for each parameter (*N*_1_*, N*_2_*, N*_3_, and selection). For each parameter, the top 25 statistics were chosen, according to the procedure in Algorithm 1. The Venn diagram captures statistics common to each subset of parameters, with notable less informative statistics shown in the lower right. Close, mid, and far represent the genomic region where the statistic was calculated. The numbers after each colon refer to the position of the statistic within its distribution or order. For the SFS statistics, it is number of minor alleles. For each region, there are 50 SFS statistics, 16 BET statistics (distribution between segregating sites), 30 IBS statistics, and 16 LD statistics.

### Regularization of the network weights

In deep learning, one hyper-parameter that should be investigated closely is the regularization parameter λ in the cost function, also called the weight decay parameter. (See `Methods` for details). If λ is set to be too high, large weights will be penalized too much, and interesting features of the data cannot be learned well. But if λ is set too low, the weights tend to display runaway behavior. Due to this balance, a validation procedure is typically used to find the right λ. In our case, the additional runtime of simulating more data and performing more training would be too computationally expensive, but we do provide a small validation study in Figure S2.

### Runtime

In terms of runtime, the vast majority is spent simulating the data. During training, most of the runtime is spent fine-tuning the deep network, which requires computing the cost function and derivatives many times. To speed up this computation, our deep learning implementation is parallelized across datasets, since each dataset adds to the cost function independently. This significantly improves the training time for deep learning, which can be run overnight on this dataset with a modest number of hidden nodes/layers. Once the training is completed, an arbitrary number of datasets can be tested more or less instantaneously. In contrast, each of the “training” datasets for ABC must be examined for *each* test dataset. This takes several weeks for a dataset of this size (which is why we tested ABC on a subset of the data), although it could be parallelized across the test datasets.

## Discussion

Using deep learning for population genetics is still in its infancy, and there are many directions of future work. One important advantage of deep learning is that it provides a way to distinguish informative summary statistics from informative ones. Here we have presented one method for extracting informative statistics given a trained deep network. Other methods are possible, and it is an open question which one would produce the best results. It would be interesting to down-sample statistics using a variety of methods, then compare the results. Overall, learning more about how statistics relate to parameters could be very useful for population genetics going forward.

The prospect of using deep learning to classify regions as neutral or selected is very appealing for subsequent demographic inference. There are other machine learning methods that perform such classification, but they are generally limited to two classes (selected or neutral). One exception is a study in humans [63], which classifies genomic regions as neutral or under positive, negative, or balancing selection. Although their approach does not jointly infer selection and demography, it would be interesting to see their method used on *Drosophila*.

We infer many hard sweeps in African *Drosophila,* which is perhaps expected given their large effective population size. However, when restricting our analysis to selected regions with high confidence, the numbers of hard sweeps and soft sweeps are comparable. It would be interesting to analyze our results in the context of a simulation study by Schrider *et al.* [54], which found that regions classified as soft sweeps are often truly the “shoulders” of hard sweeps. This possibility is worth investigating, as the signatures of soft sweeps and soft shoulders are extremely similar.

Deep learning can make optimal use of even a limited number of simulated datasets. In this vein, it would be interesting to use an approach such as ABC MCMC [39] to simulate data from a prior, then use deep learning on these simulated datasets. Alternatively, deep learning could be used to select informative statistics for a subsequent method such as ABC. Such combined approached could be a fruitful area of further exploration.

We would also like to apply deep learning to a wider variety of scenarios in populations genetics. Population structure and splits would be an examples, although these scenarios would most likely require different sets of summary statistics.

On the computer science side, deep learning has almost exclusively been used for classification, not continuous parameter inference. It would be interesting to see the type of continuous parameter inference presented here used in other fields and applications.

Finally, machine learning methods have been criticized for their “black-box” nature. In some sense they throw away a lot of the coalescent modeling that we know to be realistic, although this is included somewhat in the expert summary statistics of the data. It would be advantageous to somehow combine the strengths of coalescent theory and the strengths of machine learning to create a robust method for population genetic inference.

## Methods

### Deep learning details

In this section we provide the theory behind training deep networks. The notation in this section follows that of [42]. Let *x*^(^*^i^*^)^ be the vector of summary statistics for dataset *i*, and *y*^(^*^i^*^)^ the vector of parameters that dataset *i* was simulated under. If we have *m* such datasets, then together {(*x*^(1)^, *y*^(1)^),…, (*x*^(^*^m^*^)^, *y*^(^*^m^*^)^)} form the training data that will be used to learn the function from summary statistics to parameters. Deep neural networks are a way to express this type of complex, non-linear function. The first layer of the network is the input data, the next layers are the “hidden layers” of the network, and the final layer represents the network’s prediction of the parameters of interest.

#### Cost function for a deep network

An example deep network is shown in Fig. 6. The collection of weights between layer *ℓ* and layer *ℓ* + 1 is denoted 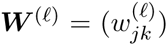, where 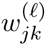 is the weight associated with the connection between node *j* in layer *ℓ* and node *k* in layer *ℓ* + 1. The biases for layer *ℓ* is denoted *b*^(^*^ℓ^*^)^. The total number of layers (including the input and output layers) is denoted *L*, and the number of hidden nodes in layer *ℓ* is denoted *u_ℓ_*. The main goal is to learn the weights that best describe the function between the inputs and the outputs.

**Figure 6.**
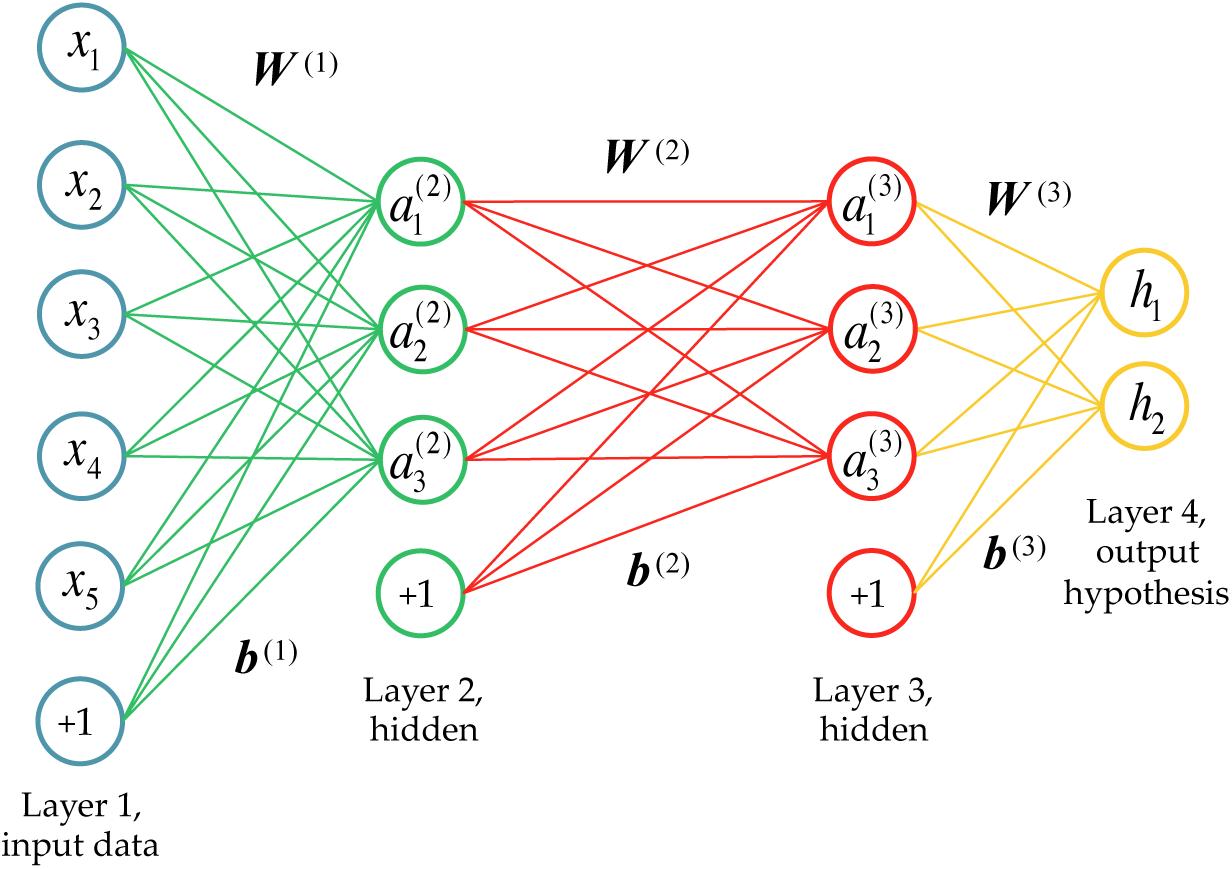
An example of a deep neural network with two hidden layers. The first layer is the input data (each dataset has 5 statistics), and the last layer predicts the 2 parameters of interest. The last node in each input layer (+1) represents the bias term. Here the number of layers *L =* 4, and the number of nodes (computational units) in each layer is *u*_1_ *= 5, u*_2_ = 3, *u*_3_ *=* 3, and *u*_4_ *=* 2 (these counts exclude the biases).

To learn this function, we first describe how the values of the hidden nodes are computed, given a trial weight vector. The value of hidden node *j* in layer *ℓ* ≥ 2 is denoted 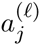, and is defined by

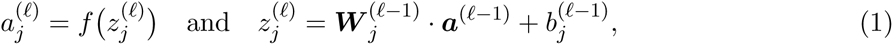

where 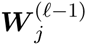 is the *j*^th^ *column* of the weight matrix ***W***^(^*^ℓ^*^−1)^ (i.e., all the weights going into node *j* of layer *ℓ* − 1), 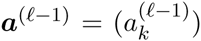 is a vector of the values of all the nodes in layer *ℓ* − 1 (with *a*^(1)^ = *x*, the input vector), and

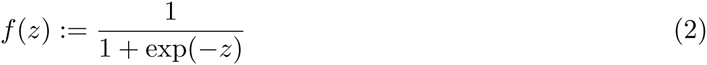

is the *activation function*. Here we use a logistic function, but other functions can be used. Another common activation function is the hyperbolic tangent function.

Hence, given the input data and a set of weights, we can *feed forward* to learn the values of all hidden nodes, and a prediction of the output parameters. These predictions are usually denoted by *hw,_b_*(*x*^(^*^i^*^)^) for our *hypothesis* for dataset *i*, based on all the weights ***W*** = (***W***^(1)^,…, ***W***^(^*^L^*^−1)^) and biases ***b*** = (***b***^(1)^,…, ***b***^(^*^L^*^−1)^). We will discuss different ways to compute the hypothesis function later on. To find the best weights, we define a loss function based on the *l*_2_-norm between this hypothesis and the true parameters. This loss function is given by

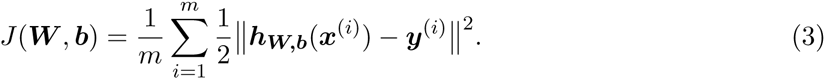

The goal of deep learning is to find the weights (***W*** and ***b***) that minimize this loss function. To efficiently find these optimal weights, we can use *backpropagation* to find the gradient. The intuition behind this approach is that once we have found the hypothesis, we then want to see how much each of the weights contributed to any differences between the hypothesis and the truth. Therefore we start at the last hidden layer, see how much each of those weights contributed, then work our way backwards, using the gradient of the previous layer to compute the gradient of the next layer. For this procedure we need to compute the partial derivatives of the cost function with respect to each weight.

Consider a single training example ***x*** with associated output ***y***. The cost for this dataset is a term in Eq. (3) and is denoted by

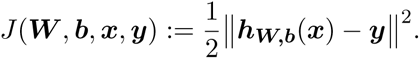

Then, define

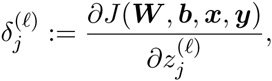

where 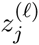 defined in Eq.(1), is the input to the activation function for node *j* in layer *ℓ.* We first consider 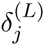 for the final layer. Noting that the *j*^th^ entry of ***hw,_b_*(*x*)** is 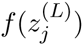, we get

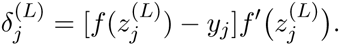

Based on this initialization, we can recursively compute all the *δ* variables:

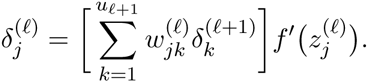

Now we can use the *δ* variables to recursively compute the partial derivatives for one dataset:

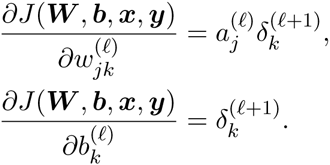

Finally, putting all the datasets together we get

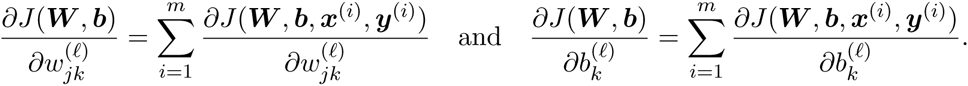

Since we can compute the derivatives using this backpropagation algorithm, we can use the LBFGS optimization routine (as implemented in [11]) to find the weights that minimize the cost function.

#### Unsupervised pretraining using autoencoders

It is possible to train a deep network by attempting to minimize the cost function described above directly, but in practice, this proved difficult due to the high-dimensionality and non-convexity of the optimization problem. Initializing the weights randomly before training resulted in poor local minima. Hinton and Salakhutdinov [23] sought to initialize the weights in a more informed way, using an unsupervised pretraining routine. Unsupervised training ignores the output (often called the “labels”) and attempts to learn as much as possible about the structure of the data on its own. PCA is an example of unsupervised learning. In Hinton and Salakhutdinov, the unsupervised pretraining step uses an *autoencoder* to try to learn the best function from the data to itself, after it has gone through a dimensionality reduction step (this can be thought of as trying to compress the data, then reconstruct it with minimal loss). Autoencoding provides a way to initialize the weights of a deep network that will ideally be close to optimal for the supervised learning step as well. See Fig. 7 for a diagram of an autoencoder.

**Figure 7.**
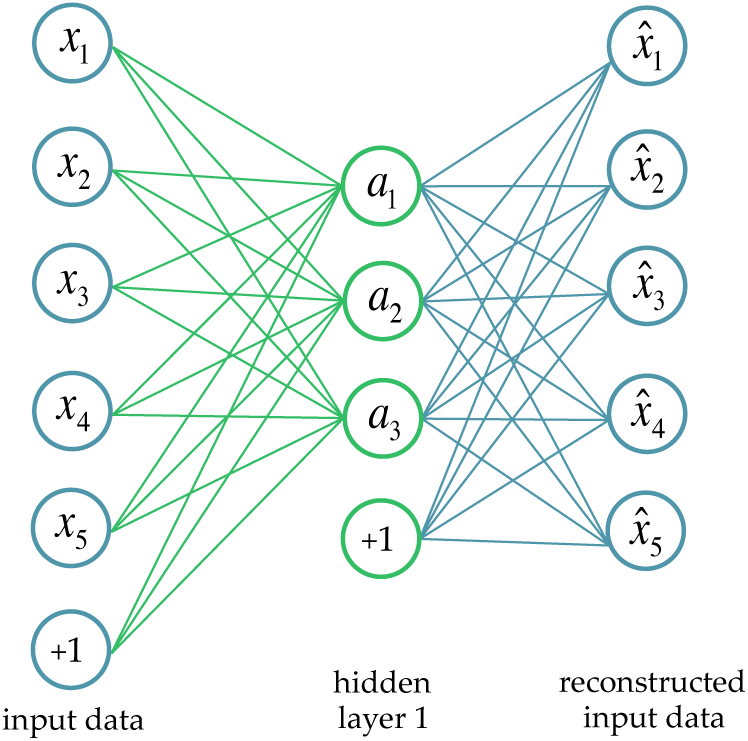
An example of an autoencoder. The input data (***x***) is projected into a (usually) lower dimension (***a***), then reconstructed 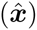 The weights of an autoencoder are optimized such that the difference between the reconstructed data and the original data is minimal.

Training an autoencoder is an optimization procedure in itself. As before, let ***W***^(1)^ be the vector of weights connecting the input *x* to the hidden layer *a,* and ***W***^(2)^ be the vector of weights connecting ***a*** to the output layer 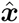, which in this case should be as close as possible to the original input data. We again typically use the logistic function shown in Eq.(2) as our activation function *f*, so we can compute the output using:

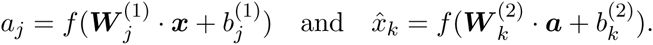

If a linear activation function is used instead of a logistic function, the hidden layer becomes the principle components of the data. This makes dimensionality reduction with an autoencoder similar in spirit to PCA, which has been used frequently in genetic analysis (see [43] for an example). However, the non-linear nature of an autoencoder has been shown to reconstruct complex data more accurately than PCA. Using backpropagation as we did before, we can minimize the following autoencoder cost function using all *m* input datasets:

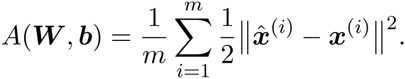

The resulting weights ***W***\*^(1)^ and ***b***\*^(1)^ will then be used to initialize the weights between the first and second layers of our deep network. The weights ***W***\*^(2)^ and ***b***\*^(2)^ are discarded. To initialize the rest of the weights, we can repeat the autoencoder procedure, but this time we will use the hidden layer 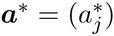, where 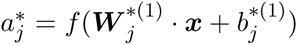, as our input data, and feed it through the next hidden layer. In this way we can use “stacked” autoencoders to initialize all the weights of the deep network. Finally, the supervised training procedure described in the previous section can be used to *fine-tune* the weights to obtain the best function from the inputs to the parameters of interest.

When the number of hidden nodes is large, we would like to constrain an autoencoder such that only a fraction of the hidden nodes are “firing” at any given time. This corresponds to the idea that only a subset of the neurons in our brains are firing at once, depending on the input stimulus. To create a similar phenomenon for an autoencoder, we create a *sparsity* constraint that ensures the activation of most of the nodes is close to 0, and the activation of a small fraction, *ρ*, of nodes is close to 1. Let 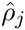 be the average activation of the hidden node *j*:

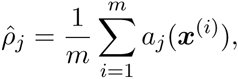

where *a_j_*(*x*^(^*^i^*^)^) is the value of the *j*^th^ hidden node when activated with dataset ***x***^(^*^i^*^)^. To ensure sparsity, we would like 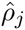 to be close to *ρ*, our desired fraction of active nodes. This can be accomplished by minimizing the KL divergence:

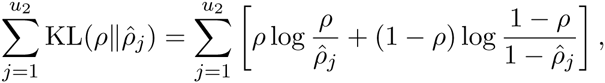

where *u*_2_ is the number of nodes in the hidden layer of the autoencoder. We multiply this term by a sparsity weight *β*.

In addition, a regularization term is included, which prevents the magnitude of the weights from becoming too large. To accomplish this, we add a penalty to the cost function that is the sum of the squares of all weights (excluding the biases), weighted by a well-chosen constant λ, which is often called the *weight decay parameter.* Including both sparsity and regularization, our final autoencoder cost becomes:

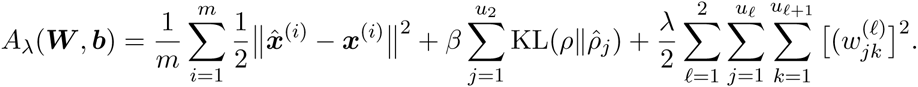

We also regularize the weights on the last layer during fine-tuning, so our deep learning cost function becomes:

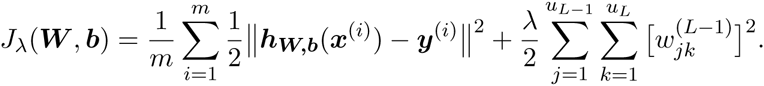

#### The final layer: parameter estimation vs. classification

In population genetics, often we want to estimate continuous parameters. To compute our hypothesis for a parameter of interest, based on a set of weights, we could use a logistic activation function, Eq.(2), as we did for the other layers. However, the logistic function is more suitable for binary classification. Instead, we use a linear activation function, so in the case of a single parameter, our hypothesis for dataset *i* becomes

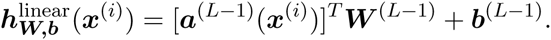

In other words, it is the dot product of the activations of the final hidden layer and the weights that connect the final hidden layer to the parameters.

For such classification results, if we had two classes, we could use logistic regression to find the probability a dataset was assigned to each class. With more than two classes, we can extend this concept and use *softmax regression* to assign a probability to each class. If we have *K* classes labeled 1,…, K, we can define our hypothesis as follows

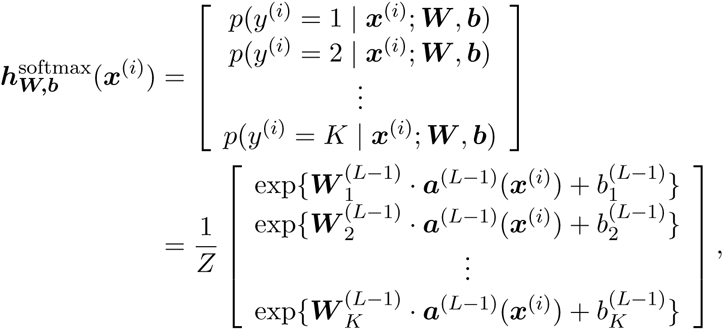

where Z is the sum of all the entries, so that our probabilities sum to 1. Using this formulation, we can define our classification cost function:

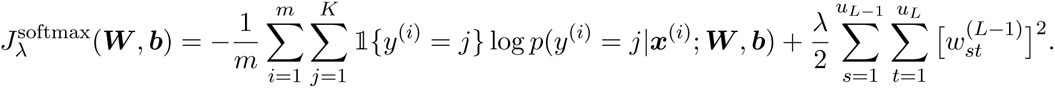

Intuitively, we can think about this cost function as making the log probability of the correct class as close to 0 as possible.

### Deep learning for demography and selection

#### Deep learning model

To modify our deep learning method to accommodate this type of inference problem, during training we have an outer-loop that changes the demography as necessary, and an inner loop that accounts for differences in selection for each region. During testing, we estimate demography and infer selection for each region separately, then average the demographies to obtain one global estimate for each genome. We also average just the estimates from the regions classified as neutral, which produces a better demographic estimate. In addition, we use these multiple estimates from each region to estimate the variance.

One final complication is that we estimate continuous parameters for the population sizes, but consider selection to be a discrete parameter. This involves a linear activation function for the population sizes and a softmax classifier for the selection parameter. A diagram of our deep learning method is shown in Fig. 8.

**Figure 8.**
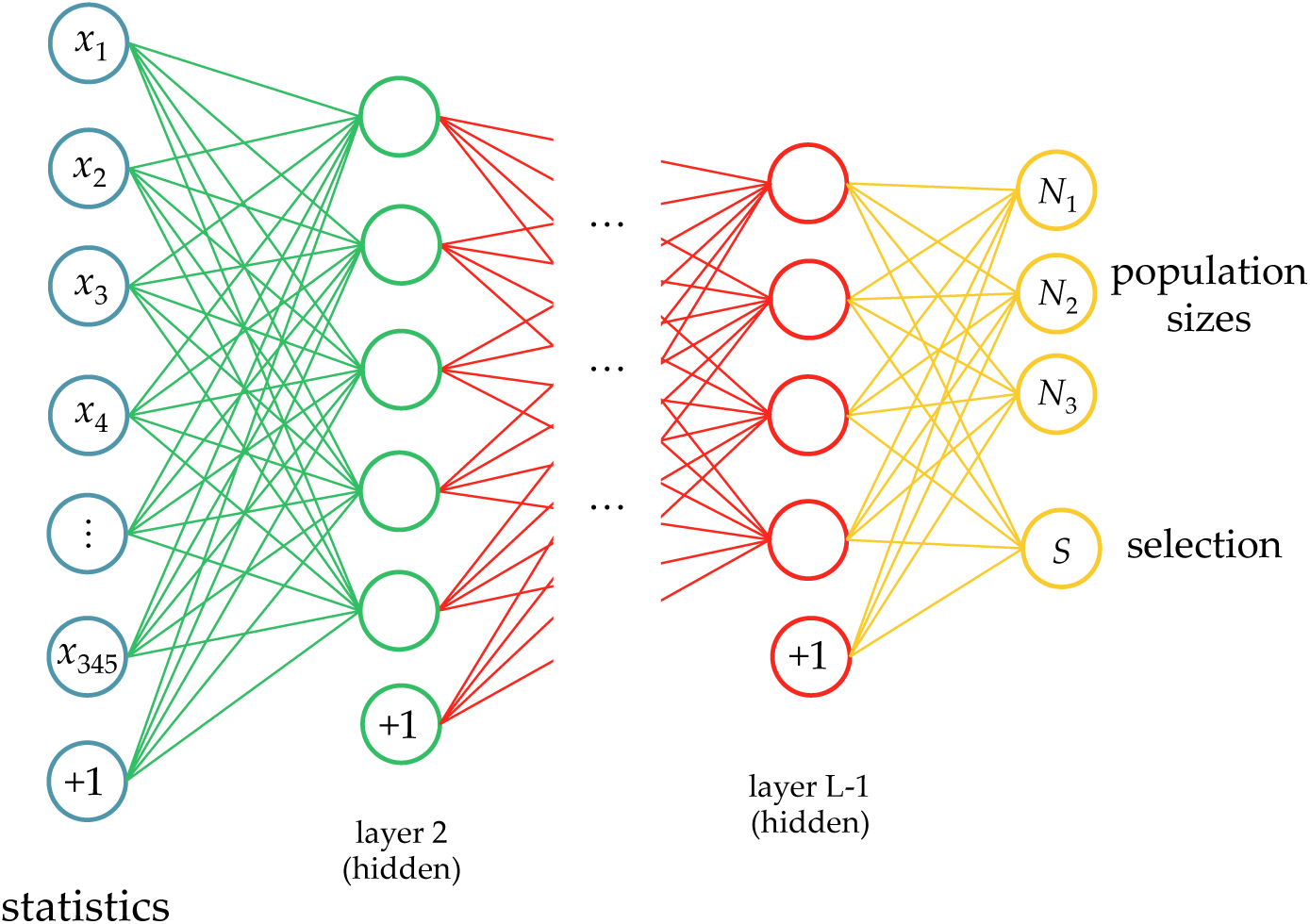
Our deep learning framework for effective population size changes and selection.

#### Analysis of informative statistics

One advantage of deep learning is that the weights of the optimal network give us an interpretable link between the summary statistics and the parameters of interest. However, from these weights, it is not immediately obvious which statistics are the most informative or “best” for a given parameter. If we only wanted to use only a small subset of the statistics, which ones would give us the best results? To answer this question, we provide a new analysis method, described in Algorithm 1.

**Algorithm 1:**
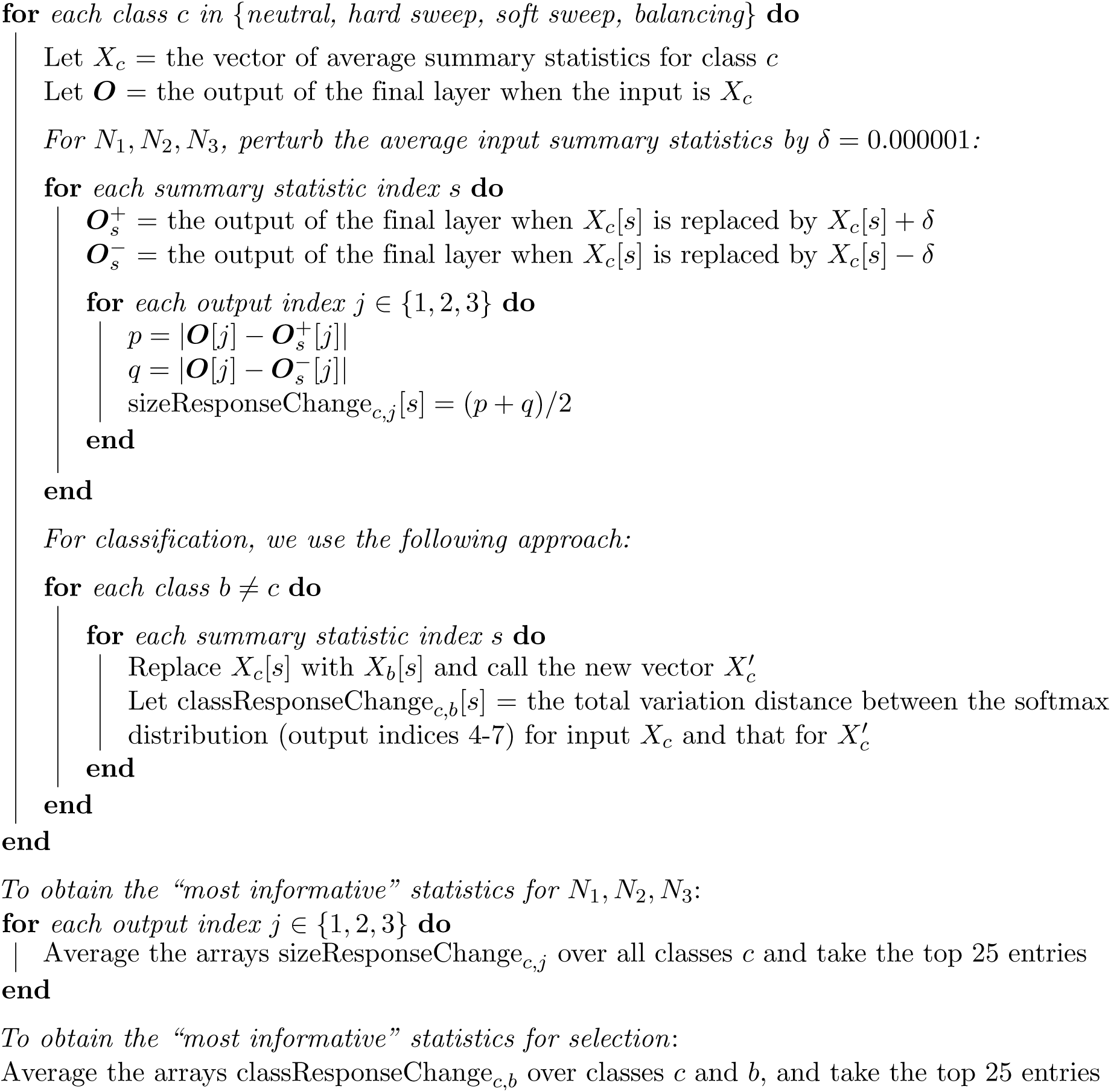
Method for finding the most informative statistics for each output parameter.

*To obtain the “most informative” statistics for selection*:

Average the arrays classResponseChange*_c_,_b_* over classes *c* and *b*, and take the top 25 entries

## Software Availability

An open-source software package that implements deep learning algorithms for population genetic inference will be made publicly available.

## Acknowledgments

We thank Andy Kern and Daniel Schrider for helpful discussion on classifying hard and soft sweeps. This research is supported in part by a National Science Foundation Graduate Research Fellowship (SS), a National Institute of Health Grant R01-GM094402 (YSS), and a Packard Fellowship for Science and Engineering (YSS).

## Supporting Information Tables and Figures

**Table S1.** Classification results for all windows, along with which genes are found in each window. (XLS)

**Table S2.** High-confidence windows, along with which genes are found in each window. (XLS)

**Figure S1.**
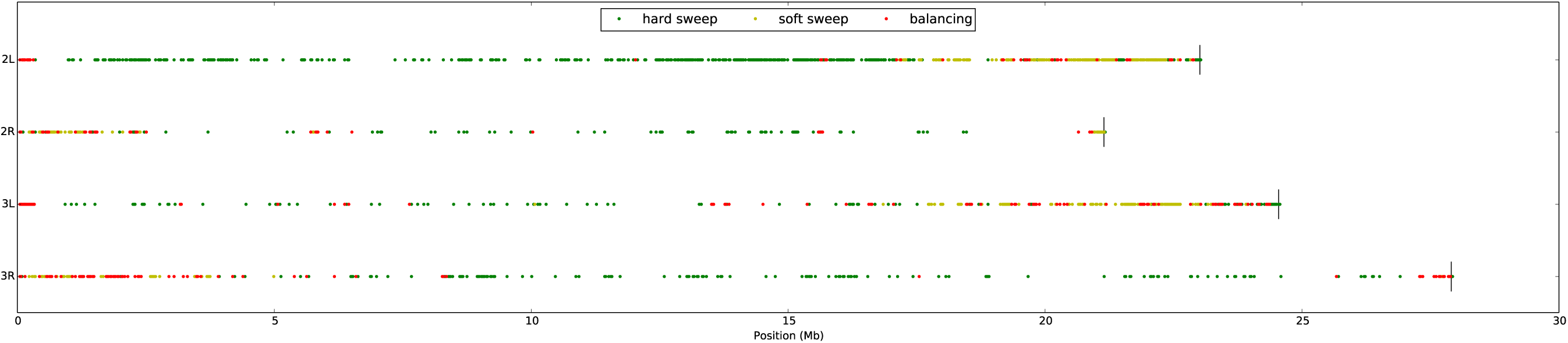
Plot of the selected regions we infer for African *Drosophila melanogaster.* Each of the 4 subplots is a chromosome arm (2L, 2R, 3L and 3R). We restricted the plot to selected sites with a probability of at least 0.95.

**Figure S2.**
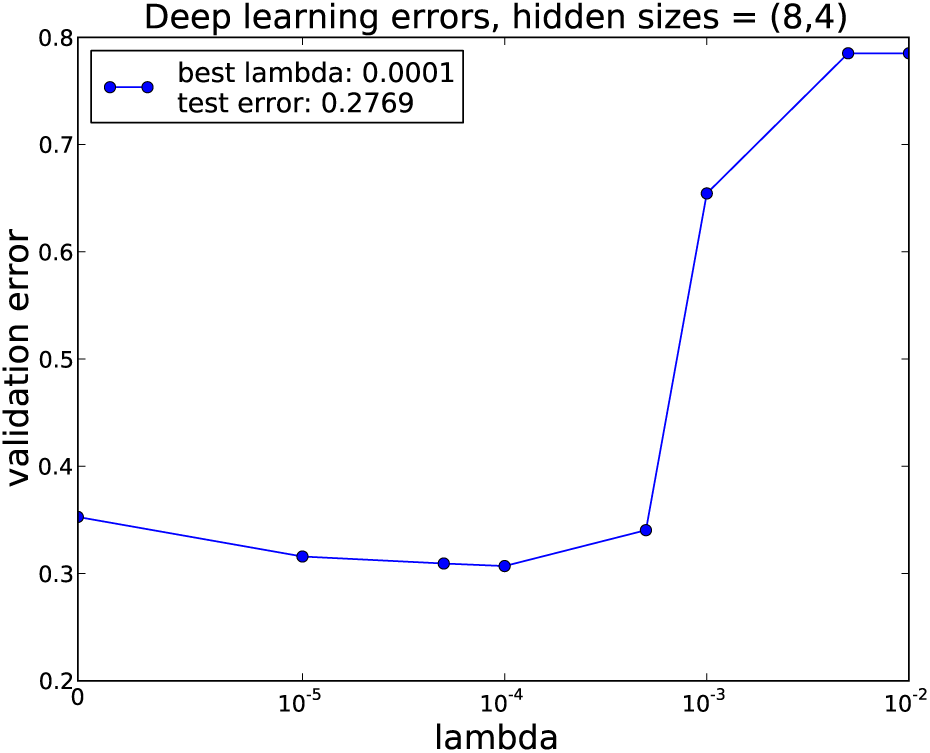
Validation procedure for a network with two hidden layers of size 8 and 4. The *x*-axis shows increasing values of λ, and the *y*-axis shows the error on the validation dataset. The curve shows a characteristic shape with low and high λ producing poorer results than an intermediate value. For these hidden layers sizes and this dataset, 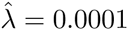 was optimal.

